# In-depth analysis of *Bacillus anthracis* 16S rRNA genes and transcripts reveals intra- and intergenomic diversity and facilitates anthrax detection

**DOI:** 10.1101/2021.08.02.454851

**Authors:** Peter Braun, Fee Zimmermann, Mathias C. Walter, Sonja Mantel, Karin Aistleitner, Inga Stürz, Gregor Grass, Kilian Stoecker

## Abstract

Analysis of 16S ribosomal RNA (rRNA) genes provides a central means of taxonomic classification of bacterial species. Based on presumed sequence identity among species of the *Bacillus cereus sensu lato* group, the 16S rRNA genes of *B. anthracis* have been considered unsuitable for diagnosis of the anthrax pathogen. With the recent identification of a single nucleotide polymorphism in some 16S rRNA gene copies, specific identification of *B. anthracis* becomes feasible. Here, we designed and evaluated a set of *in situ-, in vitro-* and *in silico-*assays to assess the yet unknown 16S-state of *B. anthracis* from different perspectives. Using a combination of digital PCR, fluorescence *in situ* hybridization, long-read genome sequencing and bioinformatics we were able to detect and quantify a unique 16S rRNA gene allele of *B. anthracis* (16S-BA-allele). This allele was found in all available *B*.*anthracis* genomes and may facilitate differentiation of the pathogen from any close relative. Bioinformatics analysis of 959 *B. anthracis* genome data-sets inferred that abundances and genomic arrangements of the 16S-BA-allele and the entire rRNA operon copy-numbers differ considerably between strains. Expression ratios of 16S-BA-alleles were proportional to the respective genomic allele copy-numbers. The findings and experimental tools presented here provide detailed insights into the intra- and intergenomic diversity of 16S rRNA genes and may pave the way for improved identification of *B. anthracis* and other pathogens with diverse rRNA operons.

## Introduction

Anthrax, caused by the spore-forming bacterium *Bacillus anthracis*, is a disease of animals but can also affect humans either through contact with infected animals and their products or as a consequence of deliberate acts of bioterrorism ^1,2^. Because of its high pathogenicity, rapid, sensitive and unambiguous identification of the pathogen is vital. However, diagnostic differentiation of *B. anthracis* from its closest relatives of the *Bacillus cereus sensu lato* group is challenging. Phenotypic properties are not species-specific and nearly identical derivatives of the anthrax virulence plasmids can also be found in related bacilli ^2^.

In spite of earlier work ^3^ rRNA gene sequences have not been deemed discriminatory for unambiguous distinction of *B. anthracis* from its closest relatives due to the lack of specific sequence variations. Recent analysis of 16S rRNA gene alleles of *B. anthracis* and relatives, however, revealed an unexpected SNP (Single Nucleotide Polymorphism) at position 1110 (position 1139 in ^4^; 1110 according to *B. anthracis* strain Ames Ancestor, NC_007530) in some of the 16S rRNA gene copies ^4^. This SNP has previously been missed, most likely because it is present only in some of the total eleven 16S rRNA gene copies ^4^. Despite the high abundance of more than 1,000 publicly available short-read genomic datasets and more than 260 genome assemblies, reliable information about sequence variations within *B. anthracis* rRNA operons is still scarce due to the limitations of short-read whole genome sequencing (WGS) and subsequent reference mapping to detect sequence variations in paralogous, multi-copy genes. Producing high-quality genomes e.g. through hybrid assemblies of long- and short-read approaches would help bridge this gap.

In this study, we validated a species-discriminatory SNP within the 16S rRNA genes of *B. anthracis* using a set of different *in situ, in vitro* and *in silico* approaches on both genomic and transcript levels. Through this work, we established new diagnostic tools for *B. anthracis* including a fluorescence *in situ* hybridization (FISH) assay and a digital PCR (dPCR) test for both genomic and transcript identification and quantification. Finally, we expanded our analysis on all short-read *B. anthracis* data sets available in the NCBI Short Read Archive (SRA) and calculated the rRNA operon copy-numbers and allele frequencies using a coverage-ratio based bioinformatics approach.

## Results

### A SNP in transcripts of 16S rRNA genes enables specific microscopic detection of *B. anthracis* by fluorescence *in situ* hybridization

Triggered by earlier data on a unique SNP position in some copies of the 16S rRNA gene of *B. anthracis* (guanine to adenine transition at position 1110) ^4^, we aimed at developing a new FISH assay for the identification of *B. anthracis*. Previous work has introduced a probe set for the FISH based identification of *B. anthracis* ^5^. Evaluation of the probe sequences revealed, however, that they are unsuitable for unambiguous *B. anthracis* identification due to unspecific probe binding ^6^. Thus, we designed a FISH probe for discriminating *B. anthracis* from all of its close relatives targeting this specific SNP in 16S rRNA genes (Probe BA_SNP_Cy3). Additionally, we developed probe BC_SNP_FAM which binds to 16S rRNA sequence found in all *B. cereus s. l*. strains, including *B. anthracis* (Supplementary Table S2). No other bacterial or archaeal 16S rRNA gene in the SILVA database had a full match for both of the newly designed probes (assessed 2021-03-01). In order to increase signal intensity and stringency ^7^ we incorporated two locked nucleic acids (LNA) in probe BA_SNP_Cy3 and one LNA in probe BC_SNP_FAM. Optimum formamide concentrations in the hybridization buffer of this FISH assay was titrated and finally set at 30% (v/v) formamide for species differentiation (Supplementary Figure S2).

For assay validation, the 16S rRNA probes were tested against a broad panel of *B. cereus s. l*. strains (Supplementary Table S1). The FISH assay allowed differentiation of *B. anthracis* from all other *B. cereus s. l*. group members. *B. anthracis* cells displayed red fluorescence Cy3-signals after hybridization of the specific 16S rRNA variation at position 1110, and green fluorescence FAM-signals resulting from hybridization to the divergent 16S rRNA featuring no *B. anthracis* specific SNP (Figure 1). No red Cy3-signals were detected in any of the non-*B. anthracis B. cereus s. l*. group strains.

**Figure 1.**
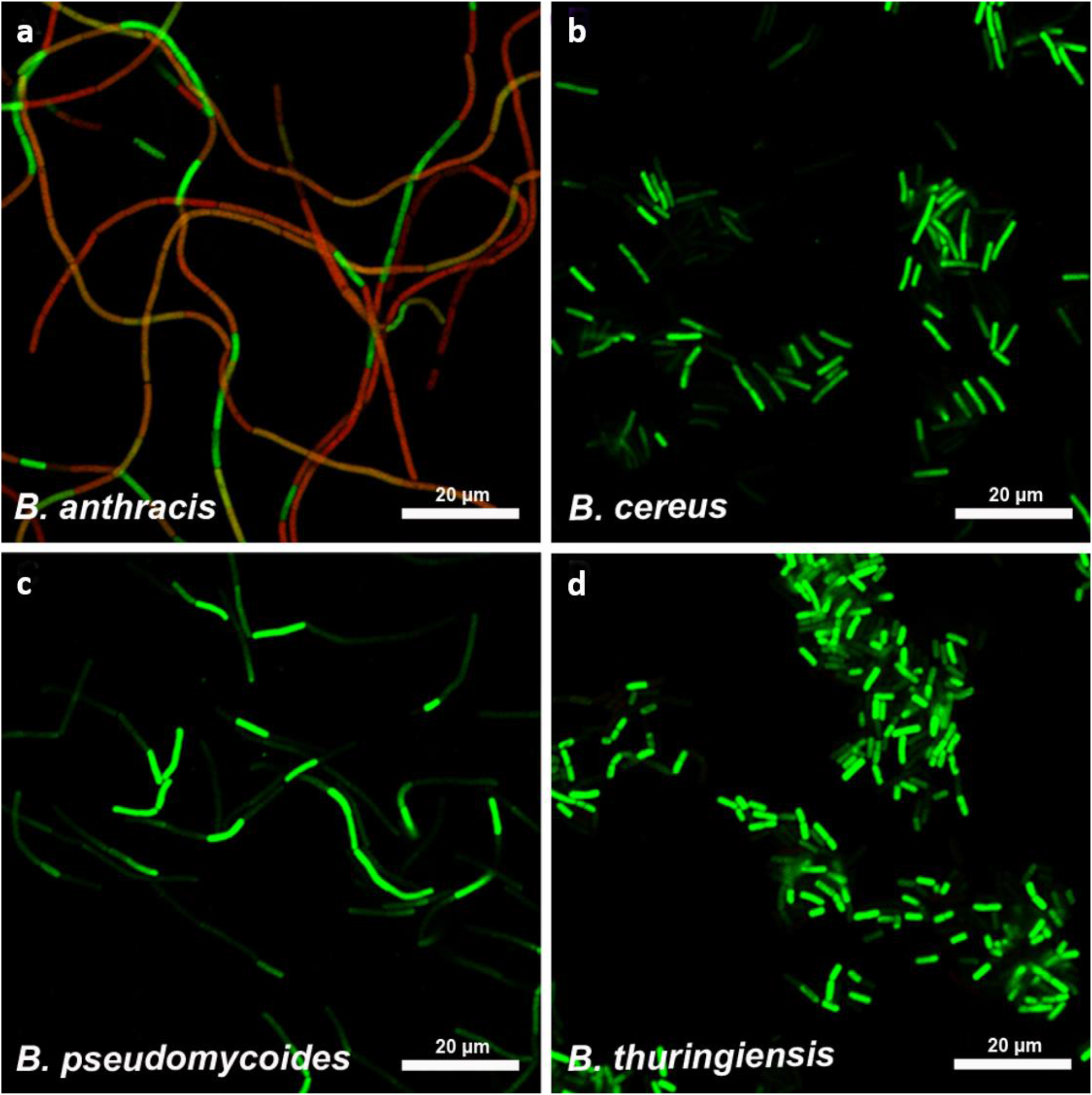
FISH-based microscopic differentiation of *B. anthracis* from other *B. cereus s. l*. group species. Representative images for *B. anthracis* (a, strain Bangladesh 28/01) *B. cereus* (b, strain ATCC 6464), *B. pseudomycoides* (c, strain WS 3119) and *B. thuringiensis* (d, strain WS 2614) are shown as overlay images of red (probe BA_SNP_Cy3 / 568nm) and green fluorescent channels (probe BC_SNP_FAM / 520nm).

While we found Cy3 FISH signals for all *B. anthracis* strains, we discovered broad variations in Cy3 fluorescence signal intensities for different cells of the same and between different *B. anthracis* strains. Even for cells of the same chain, there were individual cells showing almost uniquely either the Cy3 or the FAM signal, resulting in a mosaic-like pattern (Figure 1).Total fluorescence intensities varied between different *B. anthracis* strains from very strong Cy3 signals to the extreme cases of *B. anthracis* strains ATCC 4229 Pasteur, SA20 and A3783, for which Cy3 signals were very weak (for signal intensities see Supplementary Table S1). These findings strongly indicate that the 16S rRNA of *B. anthracis* can be used for microscopy-based specific pathogen detection. Notably, variations in fluorescence intensities suggests differences in the rRNA expression level. As these differences might be caused by a gene dose effect we decided to analyze the genomic distribution of the *B. anthracis* specific SNP in 16S rRNA genes.

### Genomic analysis of *B. anthracis* genomes reveals variations in 16S-BA-allele frequencies

We correlated FISH results with the abundance of 16S rRNA gene copies harboring the *B. anthracis* specific SNP within different *B. anthracis* genomes. Despite the significant number of *B. anthracis* genomes published, the vast majority of sequences has been generated using short-read-sequencing with subsequent mapping to the reference genome (Ames Ancestor NC_007530, ^8^). Due to multiple copies of the rRNA operons, conventional short-read-sequencing and mapping approaches do not allow for reliable detection of allele variations. During *de novo* assembly of short reads, near identical regions like rRNA operon are collapsed into one contig representing only a consensus sequence missing any minor allele variations. Thus, potential differences in allele frequencies can easily be missed. Because of mapping to the reference genome, consensus sequences always feature identical 16S rRNA allele distribution as the reference. Hence, there is a need for high-quality genomes generated by hybrid assemblies using long- and short-read-sequences for obtaining insights into the real distribution and diversity of 16S rRNA alleles in *B. anthracis* genomes.

To start meeting this need, we analyzed and compared the 16S gene sequences and locations in all available high-quality genomes of *B. anthracis* (assessed at the end of 2020) that are based on long-read-sequencing and *de novo* assembly. Figure 2 shows a schematic illustration of the genomic organization of rRNA operons including 16S, 23S and 5S ribosomal subunits as well as tRNA genes from operons *rrnC* to *H* (outlying operons A, B, I, J and K are not shown) of representative strains for different 16S rRNA genotypes (Ames Ancestor - NC_007530, ^8^; ATCC 14578 Vollum (in-house sequenced; this work; Supplementary Table S3); ATCC 4229 Pasteur - NZ_CP009476, ^9^), and closely related *B. cereus* strain ATCC 10987 - NC_003909, ^10^).

**Figure 2.**
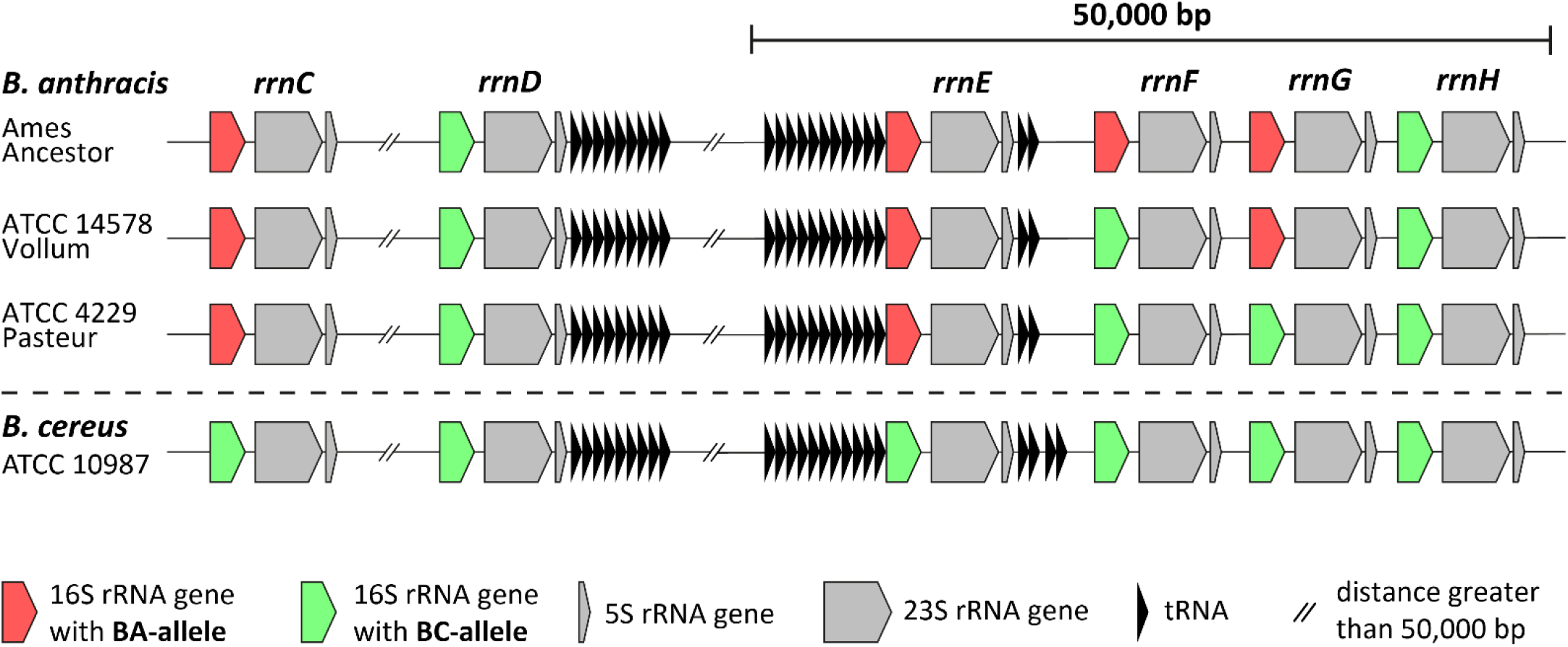
Schematic illustration of the genomic organization of rRNA operons and distribution of 16S alleles in *B. anthracis*. Depicted are the 16S, 23S, 5S ribosomal subunit, and tRNA genes from operons *rrnC* to *H* in strains Ames Ancestor, ATCC 14578 Vollum, ATCC 4229 Pasteur and *B. cereus* ATCC 10987. The 16S rRNA genes are either displayed in red for 16S-BA-alleles or in green for 16S-BC-alleles. Not shown are operons *rrnA, B, I, J* and *K* exclusively carrying the 16S-BC-allele in any strain. Distances are not to scale.

We found that all 16S rRNA gene copies featuring the *B. anthracis* specific SNP to have 100% sequence identity representing a distinct allele. For simplification, copies featuring this guanine to adenine transition at position 1110 were termed 16S-BA-(*B. anthracis*)-alleles, while all other variants lacking this transition were designated 16S-BC (*B. cereus s. l*.)-alleles.

The three *B. anthracis* strains Ames Ancestor, ATCC 14578 Vollum and ATCC 4229 Pasteur analyzed above, harbored different 16S-BA/BC-allele frequencies with 4/7, 3/8 and 2/9 copies, respectively (Figure 2). No 16S-BA-alleles were found in *B. cereus* ATCC 10987 or any other non-*B. anthracis* strain. In all three *B. anthracis* strains, rRNA operons *rrnA, B, D, H, I, J* and *K* carried 16S-BC-alleles, while for *rrnC* and *rrnE* exclusively the 16S-BA-allele was identified. Only two rRNA operons, *rrnF* and *rrnG*, were found to be variable, with strain Ames Ancestor harboring two 16S-BA-alleles and strain ATCC 4229 Pasteur only the BC-alleles for *rrnF* and *rrnG*. Strain ATCC 14578 Vollum exhibited an intermediate state with a 16S-BA-allele in *rrnG* and a BC-allele in *rrnF* (Figure 2). It is thus possible that these differences in 16S rRNA allele distributions may have caused the observed variations in *B. anthracis* specific FISH signals (Figure 1) by gene-dosage-mediated differences in rRNA transcription levels.

### A tetraplex dPCR assay enables the absolute quantification of species-specific 16S rRNA gene allele numbers in *B. anthracis*

To verify this finding and to quantify the ratios of each allele in a diverse panel of *B. anthracis* strains, we designed and tested a hydrolysis-probe-based digital PCR (dPCR) assay (Figure 3a). This assay utilized HEX (green) and FAM (blue) fluorescent dyes labelled allele-specific probes for the 16S-BC-allele and BA-allele, respectively, with both probes targeting the 1110 SNP of the 16S rRNA genes (Supplementary Table S2). In parallel, a previously published second hydrolysis-probe-based PCR assay using HEX dye was adopted for dPCR. This assay targets the *B. anthracis* specific, chromosomal *PL3* gene ^11^. Finally, a pan-*B. cereus s. l*. hydrolysis-probe-based PCR assay on the *gyrA* (gyrase gene) marker using FAM dye was designed, facilitating the detection and quantification of *B. cereus s. l*. species (including *B. anthracis*) chromosomes. In these dPCR assays, the *PL3* and *gyrA* dPCR-tests served as internal controls (for *B. anthracis* and *B. cereus s*.*l*., respectively): each positive for *B. anthracis* genomic DNA vs. negative for *PL3* and positive for *gyrA* using genomic DNA of other members of the *B. cereus s. l*. group.

**Figure 3.**
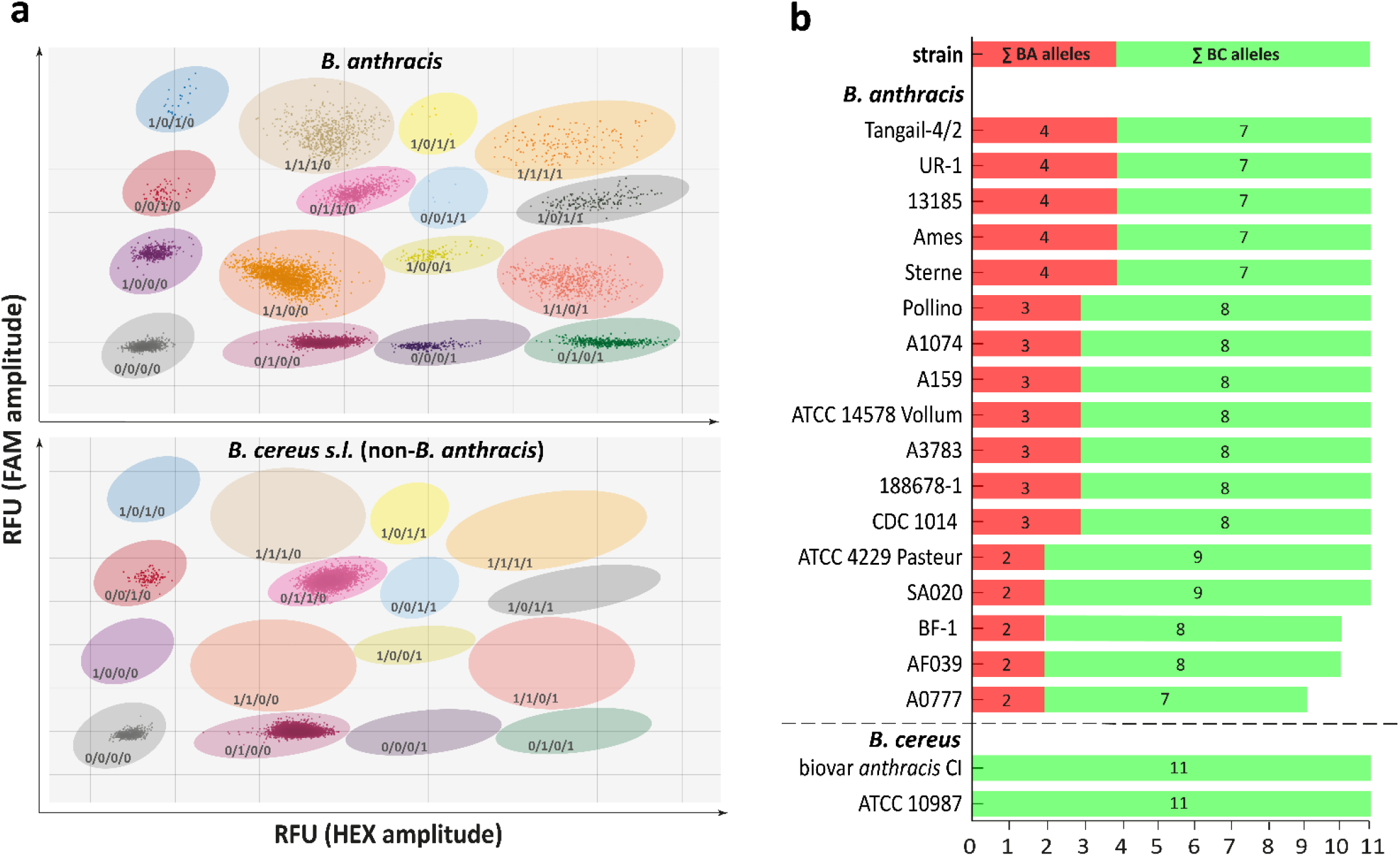
Detection and quantification of 16S rRNA gene alleles in *B. anthracis* and *B. cereus s. l*. strains. (a) typical results of a tetraplex dPCR assay using *B. anthracis* template DNA (upper panel) and DNA of a non-*B. anthracis* member of the *B. cereus s. l*. group (lower panel). With each dot representing a droplet plotted according to its FAM signal-amplitude (RFU: Relative Fluorescence Units) on the y-axis and HEX signal-amplitude on the x-axis, a total of 16 (for *B. anthracis*, upper panel) or 4 (non-*B. anthracis* members of the *B. cereus sensu lato* group, lower panel) clusters (defined by shaded areas) can be assigned to a certain dPCR marker combination of *gyrA* (FAM high signal), *PL3* (HEX high signal), 16S-BA- (FAM low signal), and 16S-BC-allele (HEX low signal). Since both the *PL3* gene and the 16S-BA-allele are exclusively found in *B. anthracis*, the 16S-BA- and 16S-BC-allele copy numbers can be calculated from the positive droplets of single copy genes (*PL3* and *gyrA*) and multi-copy 16S rRNA genes. All dPCR patterns lacking either (or both) the *PL3* gene and the 16S-BA-allele clusters represent DNA of a non-*B. anthracis* member of the *B. cereus s. l*. group. (b) Copy-numbers for 16S-BA- and 16S-BC-alleles for all *B. anthracis* strains (and *B. cereus* biovar *anthracis* CI) tested.

These four assays were combined into a single tetraplex dPCR assay. To achieve the required signal separation of the four individual dPCR reactions (on our dPCR-analysis instrument featuring only two channels, FAM and HEX), we deliberately altered the signal output levels by titrating concentrations of probes labeled with the same dye (Figure 3a). Thus, the *PL3* marker assay was tuned to produce high HEX signals vs. low HEX signals coming from 16S-BC-alleles. Likewise, the *gyrA* marker assay was set to produce high FAM signals vs. low FAM signals originating from 16S-BA-alleles. Since both *PL3* and *gyrA* are single-copy genes located on the chromosome of *B. anthracis*, these markers should result in very similar quantitative outputs when individual *B. anthracis* DNA samples are analyzed. Therefore, these markers served as internal quantification controls in this work.

A typical analysis output of this tetraplex dPCR assay is exemplified in Figure 3a. In a two-dimensional plot (FAM signal amplitude on the y-axis and HEX signal amplitude on the x-axis) of such tetraplex dPCR data, one can discriminate a specific fluorescence patterns after dPCR representing 16 clusters (when *B. anthracis* DNA was used as a template). Each of the droplets within a cluster contained a certain target combination of *gyrA, PL3*, 16S-BA-allele and/or 16S-BC-allele (for example *gyrA*^*+*^/*PL3*^*+*^/16S-BC-allele^+^/16S-BA-allele^+^ or *gyrA*^*-*^/*PL3*^*-*^ /16S-BC-allele^-^/16S-BA-allele^-^). Using template DNA originating from a non-*B. anthracis* member of the *B. cereus s. l*. group (i.e., not harboring any 16S-BA-allele), resulted in the expected formation of only four droplet clusters i.e., lacking all signals of *B. anthracis*-specific clusters containing combinations of the *PL3* marker or the 16S-BA-allele (Figure 3a).

Testing the assay on the reference strains Ames, ATCC 14578 Vollum and ATCC 4229 Pasteur we found four, three and two 16S-BA-alleles, respectively, and eleven 16S rRNA total copies per cell in all three strains. This agreed with the values determined by genomic analysis and, therefore, validated the dPCR assay being able to accurately quantify 16S rRNA alleles in *B. anthracis*.

Using the validated tetraplex dPCR assay we analyzed the same strain panel as tested by FISH (Supplementary Table S1). Similar to FISH, there was no signal for 16S-BA-alleles in the 32 non-*B. anthracis* strains of the *B. cereus s. l*. group. However, all of the 17 *B. anthracis* strains harbored at least two (up to four) copies of the 16S-BA-allele per cell (Figure 3b). The majority of *B. anthracis* strains exhibited either the genotypes 4/7 or 3/8 (16S-BA/BC-alleles; six and seven strains, respectively). These predominant genotypes, together with genotype 2/9 (strain ATCC 4229 Pasteur and strain SA020) were all found to harbor eleven rRNA operons in total, which agrees with previously determined numbers of rRNA operons in these strains. Conversely, strains A182 and BF-1 harbored only ten 16S gene copies in total (genotype 2/8). Notably, strain A0777 exhibited just nine rRNA copies, two of which contained the *B. anthracis* specific SNP (genotype 2/7)

### 16S-BA-allele frequencies and total rRNA operon copy numbers vary between different *B. anthracis* strains

In order to further confirm dPCR results and to exclude underestimation by dPCR as a possible cause of the unexpected low number of total rRNA operons in strains A182, BF-1 and A0777, we conducted a combination of long- and short-read sequencing on these and 32 additional *B. anthracis* strains (Supplementary Table S1). A mean read-length of about 15 kb generated by Nanopore sequencing combined with Illumina 2×300 bp paired-end sequencing allowed for the precise assembly of complete genomes including correct positioning of rRNA operons on chromosome. Coverage values of more than 200-fold enabled the accurate quantification of SNPs and therefore, genotypes based on 16S-BA/BC-allele distribution could be reliably determined. The results matched those obtained from dPCR, confirming the accuracy and reliability of the tetraplex assay. We found that strain A0777 lacked rRNA operons *rrnG* and *rrnH*. rRNA operon *rrnG* was not present in strains AF039, SA020 and BF-1. The genome regions downstream of the missing rRNA operons and upstream of the next rRNA operon were also absent.

In order to extend our analysis of 16S rRNA allelic states to more *B. anthracis* strains, we expanded our investigation on all publicly available short-read sequence data for *B. anthracis* generated using Illumina sequencing technology. Starting from our newly-generated high-quality hybrid assemblies, we developed a *k*-mer and coverage-ratio based tool to calculate the rRNA operon copy-numbers and allele frequencies from all SRA datasets, published until the end of 2020. These numbers of rRNA operons and 16S-BA-alleles (from short-read datasets) were identical to the long-read data of the same genomes (Supplementary Table S4). After this method validation, we analyzed 986 SRA Illumina sequenced datasets for 16S rRNA operon and BA/BC-allele distribution. After assembly and filtering, 959 genomes remained for a detailed comparison. The majority (n=735, 76.64%) contained 11 rRNA operons, 189 genomes (19.71%) harbored 10 rRNA operons and only 35 genomes (3.65%) contained 9 rRNA operons (Table 1). This ratio is comparable to that found in our initial strain-set tested with FISH and dPCR (11 copies: 82.35%, 10 copies: 11.76% and 9 copies: 5.88%). Of these 959 genomes the 16S-BA-allele distributions were: 23.04% had 2, 58.39% had 3 and 17.10% had 4 copies (Table 1), respectively. As with the rRNA operon copy-numbers, this distribution correlated with the 16S-BA-allele distribution in our strain-set analyzed by dPCR and WGS (2: 29.41%, 3: 41.17%, 4: 29.41%). Notably, a few strains were calculated to possess 1 (0.31%) or 5 (1.15%) 16S-BA-alleles. The overall diversity of 16S rRNA genotypes (BA alleles/BC alleles) was higher than in our initial strain-set (genotypes 4/7, 3/8 and 2/9). Additional major genotypes (frequency >5) obtained from SRA analysis comprised 16S-BA-/BC-allele-ratios of 2/8 and 2/7, minor genotypes were 5/6, 4/6, 4/5, 3/7, 3/6, 1/9 and 1/8, each with frequencies < 5.

**Table 1:** 16S rRNA genotypes obtained from *k*-mer based SRA analysis. Numbers of 16S-BA-alleles, overall rRNA operon numbers, and 16S rRNA genotypes resulting from these values are listed with their respective frequencies.

| 16S-BA-alleles | # of strains | # of rRNA operon copies per genome | 16S rRNA genotype [16S-BA-alleles / BC-alleles] | # of strains |
| --- | --- | --- | --- | --- |
| 1 | 3 (0.31%) | 9 | 1 / 8 | 1 (0.10%) |
|  |  | 10 | 1 / 9 | 2 (0.21%) |
| 2 | 221 (23.04%) | 9 | 2 / 7 | 28 (2.92%) |
|  |  | 10 | 2 / 8 | 116 (12.10%) |
|  |  | 11 | 2 / 9 | 77 (8.03%) |
| 3 | 560 (58.39%) | 9 | 3 / 6 | 3 (0.31%) |
|  |  | 10 | 3 / 7 | 39 (4.10%) |
|  |  | 11 | 3 / 8 | 518 (54.01%) |
| 4 | 164 (17.10%) | 9 | 4 / 5 | 3 (0.31%) |
|  |  | 10 | 4 / 6 | 32 (3.34%) |
|  |  | 11 | 4 / 7 | 129 (13.45%) |
| 5 | 11 (1.15%) | 11 | 5 / 6 | 11 (1.15%) |

Interestingly, ten of the genomes which were calculated to possess five BA-alleles are from the same originating lab and were sequenced with 100 bp single-end technique only (Supplementary Table S4). Thus, without genomic context it is hardly possible to validate the presence of a fifth 16S-BA-allele from single-end short reads. The same applies to the only other strain (BC038/2000031523) sequenced with 2×100 bp paired-end reads and a mean insert size of 520 bp. Along with three strains putatively containing a single 16S-BA-allele only, strains with five BA-alleles should be re-sequenced using long-read technology for validation.

Finally, we tested to which degree 16S rRNA genotypes fit the phylogenetic placement of strains. For this we correlated established phylogeny of *B. anthracis* based on a number of canonical SNPs ^12^ with the distribution of 16S-BA-alleles within ten major canonical SNP groups of the three branches A, B and C of *B. anthracis*. Figure S2 shows that there is limited correlation. Notably, B-branch featured a small set of genotypes besides the major 2/8 type. The few C-branch strains all had the 3/7 genotype. A-branch (comprising the majority of isolates) was the most diverse, dominantly showing the 2/9 genotype (with the exception of canSNP group Ames: 4/7). Although the 16S rRNA genotypes did not follow the established phylogeny of *B. anthracis*, the newly developed tools (tetraplex dPCR and *k-*mer based SRA analysis) might still be harnessed as an alternative typing system for *B. anthracis* strains.

### Expression of 16S-BA-alleles is proportional to gene copy-number

Varying ratios of 16S-BA/BC-alleles constitute possible explanations for differences in FISH signals of cells of diverse *B. anthracis* strains (compare Figure 1). Indeed, we found a significant correlation between 16S-BA/BC-allele ratios in sequenced genomes and mean intensities of the Cy3 FISH signals targeting the16S-BA-allele (tested with the cor.test function in R, Pearson’s r=0.61, p-value=0.009), confirming this assumption.

In order to investigate whether the 16S-BA-alleles are differentially expressed throughout different growth phases of *B. anthracis* we quantified 16S rRNA from growth experiments (Figure 4). For this, culture samples of *B. anthracis* strains Sterne, CDC 1014 and Pasteur ATCC 4229 representing three major 16S-BA/BC-allele genotypes 4/7, 3/8 and 2/9, respectively, were taken for total RNA extraction at several time points during lag, log and stationary growth phase. To compare rRNA levels with FISH signals, we also took parallel samples from six of these time points for FISH analysis. By a one-step reverse transcription duplex dPCR, the two 16S allele targets were interrogated for the expression ratios of the 16S-BA-*vis-à-vis* the BC-alleles. *B. anthracis* RNA yielded four clusters of droplets in 2D analysis plots, namely 16S-BC-allele^-^/16S-BA-allele^-^, 16S-BC-allele^+^/16S-BA-allele^-^, 16S-BC-allele^-^/16S-BA-allele^+^ and 16S-BC-allele^+^/16S-BA-allele^+^ (Figure 4). RNA of other *B. cereus s. l*. strains produced only two cluster types lacking 16S-BC-allele^-^/16S-BA-allele^+^ and 16S-BC-allele^+^/16S-BA-allele^+^. Absolute quantification of the two initial target concentrations of 16S-BA-alleles/BC-alleles in samples from growth cultures made it possible to determine their ratios representing the expression levels of the 16S-BA-alleles relative to those of 16S-BC-alleles (Figure 4). Notably, 16S-BA/BC-allele rRNA ratios varied during growth and showed similar expression patterns in all three tested *B. anthracis* strains. Starting from a relatively low 16S-BA/BC-allele ratio in early log phase, the fraction of 16S-BA-allele expression increased in early log-phase and decreased in mid log-phase with a final increase towards the stationary phase. While shifts in 16S-BA/BC-allele expression patterns in these strains were similar, differences were observed in numerical expression ratios. *B. anthracis* Sterne showed the highest 16S-BA/BC-allele expression ratio ranging from 0.44 (early exponential phase) up to 0.75 (stationary phase), compared to CDC 1014 with 0.36 to 0.69 and Pasteur ATCC 4229 with 0.22 to 0.58, which was found to have the lowest 16S-BA-allele expression in all growth phases. The largest differences in expression levels between all strains were observed in late log phase (Figure 4). The observed diverging levels of 16S-BA-allele expression in the three tested strains can easily be explained by the different numbers of 16S-BA-allele copies per genome (2, 3 or 4). Nevertheless, the proportion of 16S-BA-allele rRNA in late-exponential *B. anthracis* cells is quite disproportionate. If all rRNA operons were transcribed at a constant and equal rate, one would expect a ratio of 0.22 (Pasteur 2/9), 0.38 (CDC 3/8), and 0.57 (Sterne 4/7). Instead, we measured ratios, which correlate to a 1.57-(Pasteur), 1.46-(CDC), and 1.09-(Sterne) fold 16S-BA-allele over-representation on average throughout all growth phases and up to 2.59 - (Pasteur), 1.83-(CDC), and 1.32- (Sterne) fold in stationary phase.

**Figure 4.**
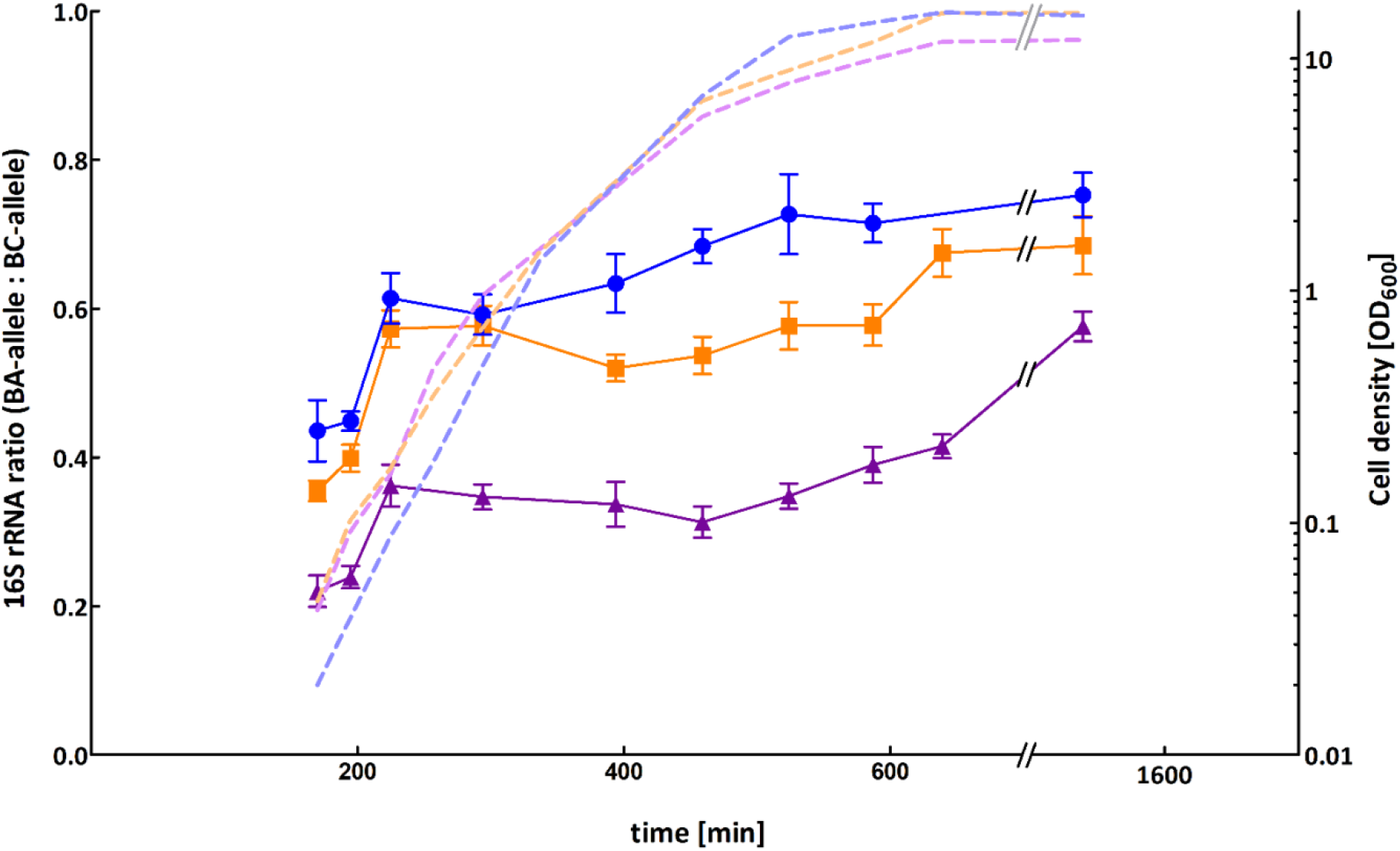
Expression ratios of 16S-BA- and –BC-alleles in three different *B. anthracis* strains at different growth phases. Expression level ratios of 16S-BA-alleles relative to 16S-BC-alleles were calculated from absolute target concentrations obtained by RT-dPCR. Values were plotted against time-points of each sample taken during growth from early exponential to stationary phase for *B. anthracis* Sterne (blue), CDC 1014 (orange) and Pasteur ATCC 4229 (purple) representing three major 16S rRNA genotypes (BA/BC) 4/7, 3/8 and 2/9, respectively. Error bars indicate the Poisson 95 % confidence intervals for each copy-number ratio. Dotted lines depict cell densities over time.

The shift towards elevated expression of the 16S-BA-allele genes over time was not significantly reflected in FISH signal intensities, possibly due to the general decrease of FISH signals over time. However, if cells were sampled and fixed at identical time points, 16S-BA/BC-allele ratios were always highest for *B. anthracis* Sterne and lowest for *B. anthracis* Pasteur, which reflects their 16S-BA/BC-allele ratios on the genomic and transcript levels (Figure 5). Also, sampled across all time points, 16S-BA/BC-allele FISH signal ratios correlated well with allele distributions in the three different strains (ANOVA in R, p= 0.0002, Figure 5).

**Figure 5.**
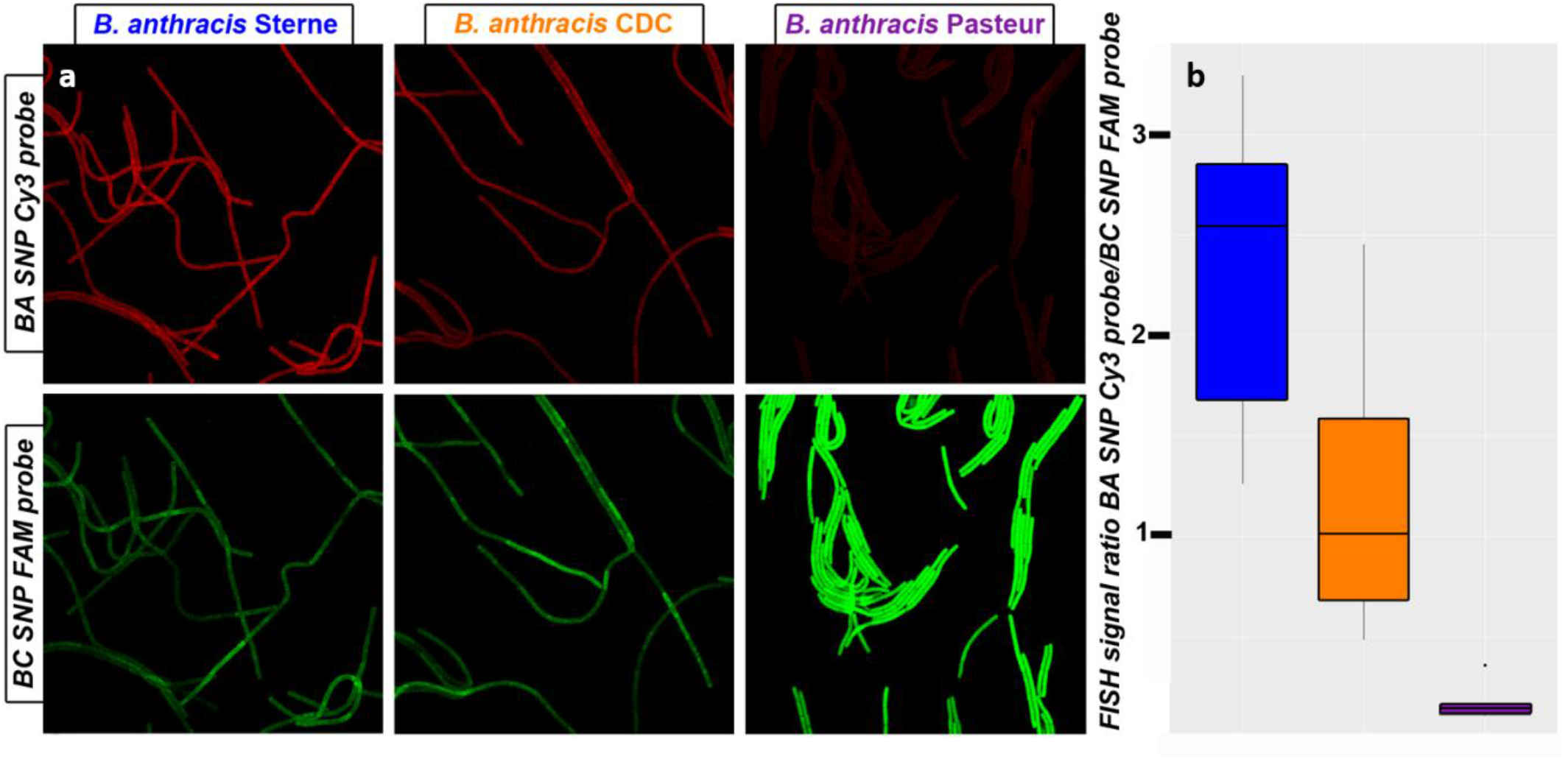
FISH of *B. anthracis* strains harboring different numbers of 16S-BA-alleles. (a) Representative FISH images showing signal intensities of *B. anthracis* strains with diverging genomic 16S (BA/BC) allele profiles (Sterne 4/7), CDC 1014 (3/8) and Pasteur ATCC 4229 (2/9). Samples were taken and processed after 460 min of continuous growth. (b) Boxplot of BA_SNP_Cy3 and BC_SNP_FAM FISH signal ratios across all sampled time points for *B. anthracis* Sterne (blue), CDC 1014 (orange) and Pasteur ATCC 4229 (purple).

## Discussion

Using a combination of newly developed *in situ, in vitro* and *in silico* approaches, we unraveled the elusive heterogeneity of 16S rRNA genes in the biothreat agent *B. anthracis*. Results consistently delineate the organism’s intragenomic diversity of 16S rRNA genes, their differential expression across growth phases and their intergenomic heterogeneity in publicly available and newly sequenced genomes. Intragenomic micro-diversity within 16S rRNA genes has long been known from other species ^13,14^ and was found to increase with higher copy-numbers of rRNA operons ^15^. Thus, the species-wide intra- and inter-genomic micro-diversity related to SNP 1110 in the 9 to 11 copies of the 16S rRNA gene of *B. anthracis* is not totally unsurprising ^3,4^. Whereas some of such polymorphic sites are associated with a distinct phenotypic trait (e.g. stress resistance)^16,17^ the functional assignment for the majority of these sequence variations (including those in *B. anthracis* 16S rRNA genes) remains elusive.

Though discovered before using Sanger sequencing ^4^, the specific SNP in the 16S rRNA genes of *B. anthracis*, was disregarded despite the availability of numerous published genomes. Generally, sequence variations in multi-copy genes such as 16S rRNA genes can hardly be detected when relying on conventional short-read WGS and subsequent reference mapping ^18^, which was used to generate the majority of publicly available *B. anthracis* whole genomes sequences. SNP calling in different rRNA operons or other paralogous genes gives ambiguous results since assemblers tend to interpret low frequent sequence variations as sequencing errors and correct them prior assembly ^19^. Even if detected, distances of the SNP to unique flanking regions up- and downstream of the multi-copy gene may be >1000 bases and thus, larger than typical library fragment sizes of 500–800 bases. In such cases, chromosomal locations of SNPs cannot be reconstructed. Instead, all rRNA gene related reads are assembled into one contig with diverse fringes ^20^. The average read length of Nanopore sequencing is typically larger than 5 kb and can therefore cover complete rRNA operons. Thus, any unique SNP occurring in a single or a few rRNA gene alleles can be precisely allocated to a specific chromosome position, especially when combined with short-read sequencing and hybrid assembly as used here. Therefore, the challenges described above will become rather minor for future genomic analysis of *B. anthracis*. Such work is facilitated by the additional 33 complete high-quality genomes we have contributed here. These genomes cover all three major phylogenetic lineages (canSNP groups), all *bona fide* 16S-BA-allele frequencies (2, 3, and 4) as well as all known rRNA operon copy numbers.

On the *B. anthracis* chromosome, the 16S rRNA operons *rrnE, F, G* and *H*, are located in close proximity to each other with only 15.8, 8.5 and 5.2 kb in-between, respectively (forming a genomic region with a high density of four 16S rRNA operons within less than 50 kb). Conversely, the other 16S rRNA genes are rather dispersed with distances greater than 50 kb in-between. The 16S-BA-allele is present in operons *rrnC* and *rrnE* in all strains analyzed with long-read WGS while *rrnF* and *rrnG* seem to be variable. Since the four operons *rrnE, F, G* and *H* are relatively close to each other in the *B. anthracis* chromosome, homologous recombination and gene duplications might be the reason for this allelic variation. Also, compared to all other 16S rRNA alleles on the *B. anthracis* genome, the 16S rRNA copies in this region (*rrnE* – *rrnH*) seem to differ from each other only in SNP position 1110. This finding promotes the explanation that the 16S rRNA copies in this region of high rRNA operon density are subjected to an increased recombination-rate between alleles with and without *B. anthracis* specific SNP 1110. This notion is also supported by the fact that only operons *rrnG* and *H* seem to be affected by deletion events in all strains analyzed by long-read WGS. The alternative explanation, horizontal gene transfer of a divergent allele, seems unlikely. We were unable to identify any 16S rRNA gene in public databases matching the 16S-BA-allele outside *B. anthracis*.

Recombination and deletion events in 16S rRNA operons of *B. anthracis* do occur as evidenced by a study on bacitracin resistance. Two deletion events, DelFG and DelGH, were described which caused elimination of gene-clusters between rRNA operons *rrnF* and *G* and *G* and *H*, respectively ^18^. These DelFG- and DelGH-events describe a possible origin of *B. anthracis* strains with ten 16S rRNA gene copies, i.e. 21% of all strains (Table 1). Random gene duplication and gene elimination by recombination might also explain another observation: the newly defined 16S rRNA genotypes did not convincingly reflect the established *B. anthracis* phylogeny (Supplementary Figure S1). Instead, some 16S rRNA genotypes seem to be dominant yet not exclusive in separate branches, e.g. 2/8 copies in B-branch or 3/8 in A-branch (Supplementary Figure S1).

The recognition of intra- and intergenomic 16S rRNA allele diversity in *B. anthracis* opens possibilities to harness unique SNPs in 16S rRNA gene alleles and their transcripts. This finding strongly highlights the great potential of such genomic variations for both identification of *B. anthracis* and for diagnostics of anthrax disease. This approach is probably also applicable to other pathogens which are otherwise difficult to discriminate from their less notorious relatives.

## Materials and Methods

### Cultivation of bacteria

The cultivation of the virulent *B. anthracis* strains was performed in a biosafety level 3 laboratory (BSL3). All *Bacillus* strains were cultivated overnight on Columbia blood agar plates (containing 5 % sheep blood, Becton Dickinson, Heidelberg, Germany) at 37°C. For isolation of DNA, a 1 µl loop of colonies was transferred to a 2 mL screw cap microfuge tube, inactivated with 2 % Terralin PAA (Schülke&Mayr GmbH, Norderstedt, Germany) for 30 min and washed three times with phosphate-buffered saline (PBS) as described previously ^21^.

For FISH, 50 ml centrifuge tubes containing 5 ml of Tryptic Soy Broth (TSB, Merck KGaA, Darmstadt, Germany) were inoculated with one colony from an overnight culture (see above) and incubated at 37° C with shaking at 150 rpm. After four hours of growth, bacteria were pelleted by centrifugation at 5,000 x g for 10 min, washed with PBS, and fixated with 3 ml 4% (v/v) formaldehyde for one hour at ambient temperature. After fixation, cells were washed three times with PBS, resuspended in a 1 : 1 mixture of absolute ethanol and PBS and stored at −20 °C until further use. To ensure sterility 1/10 of the inactivated material was incubated in thioglycolate-medium (Merck KGaA, Darmstadt, Germany) for seven days without growth before material was taken out of the BSL3 laboratory.

For growth phase analysis, 1 ml of overnight cultures of attenuated *B. anthracis* (Sterne, CDC 1014 and ATCC 4229 Pasteur) in TSB was used to inoculate 100 ml of fresh TSB in 1 L baffled flasks and incubated at 37°C with shaking at 100 rpm. Every 30 min, turbidity was measured as OD_600_ and 1 ml samples were taken for FISH and RNA isolation, respectively. After pelleting by centrifugation samples for RNA isolation were resuspended and inactivated using 2% Terralin PAA for 30 min and washed three times with PBS. FISH samples were treated as described above.

### Design of Primers and Probes

Primers and probes were designed using Geneious 10.1.3 (Biomatters, Auckland, New Zealand) and numerous probe variations were tested to identify the best combination and number of locked nucleic acids for differentiation of *B. anthracis* and the other *B. cereus s. l*. group species based on the SNP (pos. 1110) detected previously ^4^. The final probes for FISH included two and one locked nucleic acid while dPCR probes contained 5 and 6 for the *B. anthracis* (BA) and the *B. cereus s*-l- (BC) probe, respectively (Supplementary Table S2). For sequences of positive (EUB338 ^22^) and negative (nonEUB, ^23^) control probes for FISH, see Supplementary Table S2. Primers as well as probes labeled with 6-carboxyfluorescein (6-FAM), hexachlorofluorescein (HEX), indocarbocyanine (Cy3) or indodicarbocyanine (Cy5) were purchased commercially (TIB Molbiol, Berlin, Germany).

To determine the ideal formamide concentration for the FISH hybridization buffer, the fluorescence signals of probe BA_SNP_Cy3 and probe BC_SNP_FAM were assessed with *B. anthracis* Sterne and *B. cereus* ATCC 10987 at different formamide concentrations (0, 10, 20, 25, 30, 35, 40, 45, 50% FA concentration in the hybridization buffer) as described elsewhere ^24^. Hybridization at 30% formamide was determined to be ideal for differentiation of *B. anthracis* and *B. cereus s. l*. group (Supplementary Figure S3).

### Fluorescence in situ Hybridization and Image Processing

FISH was carried out as described elsewhere ^24^. A positive-control probe targeting eubacteria (EUB338, ^22^) and a nonsense probe targeting no known bacterial species (nonEUB, ^23^) as a control for unspecific probe binding were included in each hybridization experiment. Briefly, 2 µl of fixed cells were spotted on teflon coated slides (Marienfeld, Lauda-Königshofen, Germany) and dried at 46°C. Then, cells were permeabilized using 10 ml of 15mg/ml lysozyme (Merck KGaA, Darmstadt, Germany, Cat.Nr. 62970) per well at 46°C for 12 min. After dehydration in an ascending ethanol series (50, 80, 96% (v/v) ethanol) cells were covered with 10 µl hybridization buffer (0.9 M NaCl, 20 mM Tris-HCl (pH 8.0), 0.01% SDS, 30% formamide) with probes at a concentration of 10 µM and incubated in a humid chamber in the dark at 46°C for 1.5 h. Slides were washed in 50 ml pre-warmed washing buffer (0.1 M NaCl, 20 mM Tris-HCl (pH 8.0), 5 mM EDTA (pH 8.0)), for 10 min at 48°C in a water bath. Finally, slides were dipped in ice-cold ddH_2_O and carefully dried with compressed air. For each strain, FISH was performed in duplicate and two pictures were taken per well, so that the resulting fluorescence intensity was the mean of four images. To increase accuracy in the growth curve assay, five pictures were taken per well, so that the resulting fluorescence intensity was the mean of ten images. All images were recorded with a confocal laser-scanning microscope (LSM 710, Zeiss, Jena, Germany). Excitations for FAM, Cy3 and Cy5 were at 490, 560 and 630 nm respectively. Emission was measured within the following ranges: FAM: 493-552 nm, Cy3: 561-630 nm and Cy5: 638-724 nm. Images were processed with Daime ^25^, using the area of the EUB signal as a mask to measure average fluorescence intensity for BA_SNP_Cy3 and BC_SNP_FAM: The EUB images were segmented and unspecific fluorescence excluded with default threshold settings and this object layer was transferred to BA_SNP_Cy3 and BC_SNP_FAM images.

### Isolation of nucleic acids

DNA isolation from inactivated cells was carried out using MasterPure™ Gram Positive DNA Purification Kit (Lucigen, Middleton, WI, USA) according to the manufacturer’s protocol. DNA samples were quantified using the Qubit dsDNA HS Assay Kit protocol (Thermo Scientific, Dreieich, Germany). For RNA isolation from inactivated cells, RNeasy Protect Bacteria Mini Kit (Qiagen, Hilden, Germany) was used according to the supplier’s protocol for enzymatic lysis and proteinase K digestion of bacteria. In order to eliminate residual DNA, RNA samples were purified twice using RNeasy MinElute Cleanup Kit (Qiagen, Hilden, Germany) and quantified using the Qubit RNA HS Assay Kit protocol (Thermo Scientific, Dreieich, Germany). The absence of DNA in the final RNA preparation was verified by conducting PCR on marker *dhp61* ^26^ with negative results.

### Tetraplex droplet digital PCR assay for quantification of 16S rRNA gene alleles

Digital PCR (dPCR) allows for absolute quantification of DNA or RNA template concentrations ^27^. For 16S rRNA gene analysis the 20 μl dPCR pre-droplet mix consisted of 10 μl dPCR Supermix for Probes (Bio-Rad Laboratories, Munich, Germany), 1 µl 20x 16S SNP Primer mix (final concentrations 900 nM), 0.6 µl of 20x mix of 16S SNP BC probe (final concentration 150 nM), 0.6 µl of 20x mix of 16SSNP BA probe (final concentration 150 nM), 0.9 µl of 20x PL3 primer/probe mix (final concentrations: probe 225 nM, primers 810 nM), 1.5 µl of GyrA 20x primer/probe mix (final concentrations: probe 375 nM, primers 1350 nM), 4.4 µl of nuclease free water (Qiagen, Hilden, Germany) and 1 µl of template DNA freshly diluted to a concentration of 0.05 ng/µl. To ensure independent segregation of the 16S rRNA gene copies from the bacterial chromosome and the reference genes into droplets, template DNA was digested (no cut sites within 16S rRNA genes) prior to dPCR by BsiWI-HF, BsrGI-HF and HindIII-HF (New England Biolabs GmbH, Frankfurt am Main, Germany) in 1 x Cutsmart buffer (New England Biolabs GmbH, Frankfurt am Main, Germany) for 60 min and then the enzymes heat inactivated at 80°C for 20 min according to the manufacturer’s protocol. Partitioning of the reaction mixture into up to 20,000 individual droplets was achieved using a QX200 dPCR droplet generator (Bio-Rad Laboratories, Munich, Germany). A two-step PCR-reaction was performed on a Mastercycler Pro instrument (Eppendorf, Wesseling-Berzdorf, Germany) with the following settings: one DNA-polymerase activation step at 95°C for 10 min was followed by 40 cycles of denaturation at 94°C for 30 s and annealing/extension at 58°C for 1 min. Final Finally enzyme inactivation was performed at 98°C for 10 min before the samples were cooled down and held at 4°C. All steps were carried out with a temperature ramp rate of 2°C/s. After completion droplets were analyzed using the QX100 Droplet Reader (Bio-Rad) and absolute concentrations for each target were quantified using Poisson statistics as implemented in the Quantasoft Pro Software (Bio-Rad Laboratories, Munich, Germany).

Then, the absolute concentrations of *PL3* and *gyrA* were compared. To ensure assay integrity samples with a deviation range greater than 10% within the two markers were excluded and had to be repeated. If deviation was below 10% both targets were set as a reference with a copy-number of one. The software then automatically takes the mean concentration of both references to calculate the copy-numbers of BC and BA alleles. According to the recommendations provided by ^28^, all samples with copy-numbers between 0.35 and 0.65 deviating from an integer number or with a confidence interval greater than 1 were excluded from analysis and were repeated. All valid runs were rounded to the next integer number.

### Duplex one-step reverse transcription dPCR to compare expression levels of 16S BC- and 16S-BA-allele

The 20 μl RT-dPCR reaction mixture consisted of 5 μl One-Step RT-dPCR Advanced Supermix for Probes (Bio-Rad, Laboratories, Munich, Germany), 2 µl of Reverse Transcriptase (Bio-Rad, final concentration 20 U/µl), 0.6 µl of DTT (Bio-Rad, Laboratories, Munich, Germany; final concentration 10 nM), 1.5 µl 20x 16S SNP Primer mix (final concentrations 1350 nM), 1.5 µl of 20x mix of 16S SNP BC probe (final concentration 375 nM), 1.5 µl of 20x mix of 16S SNP BA probe (final concentration 375 nM), 6.9 µl of nuclease free water (Qiagen, Hilden, Germany) and 1 µl of template RNA. Reverse transcription was achieved within droplets prior to dPCR. Partitioning of the reaction mixture into up to 20,000 droplets was carried out using a QX200 dPCR droplet generator (Bio-Rad, Laboratories) and PCR was performed on the Mastercycler Pro (Eppendorf, Wesseling-Berzdorf, Germany) with the following settings: The initial reverse transcription step was performed at 48°C for 60 min. An enzyme activation step at 95°C was carried out for 10 min followed by 40 cycles of a two-step program of denaturation at 94°C for 30 s and annealing/extension at 58°C for 1 min. Final enzyme inactivation was performed at 98°C for 10 min before the samples were cooled down and held at 4°C. All steps were carried out with a temperature ramp rate of 2°C/s. After completion, droplets were analyzed using the QX100 Droplet Reader (Bio-Rad, Laboratories, Munich, Germany) and results were quantified with the Quantasoft Pro Software (Bio-Rad, Laboratories).

### Library preparation, sequencing and assembly of genomes

The libraries for the Illumina sequencing were prepared using the NEBNext® Ultra™ II FS DNA Library Prep Kit for Illumina (New England BioLabs GmbH, Frankfurt am Main, Germany) according to the protocol for large fragment sizes >550 bp but with a minimal fragmentation time of only 30 s. Afterwards, libraries were pooled equimolarly and sequenced on an Illumina MiSeq device (Illumina Inc., San Diego, CA, U.S.A.) using the MiSeq Reagent Kit v3 (2×300 bp).

The libraries for the nanopore sequencing were prepared using the Ligation Sequencing Kit SQK-LSK109 (Oxford Nanopore Technologies, Oxford, U.K.) combined with the Native Barcoding Expansion EXP-NBD104 and sequenced as one pool on a MinION flowcell FLO-MIN106D (Type R9.4.1; Oxford Nanopore Technologies, Oxford, U.K.) for 48 h. Basecalling and demultiplexing was done separately using Guppy v3.2.10 (Oxford Nanopore Technologies, Oxford, U.K.) with the high accuracy basecalling model. Quality (≥10 q) and length (≥1,000 bp) filtering was done using Filtlong version 0.2.0 (https://github.com/rrwick/Filtlong).

Hybrid assemblies were constructed in two stages. First, nanopore reads were assembled using Flye version 2.7 ^29^ with default parameters and two iterations of polishing. Second, Illumina reads were assembled together with the nanopore raw reads and the nanopore assembly as trusted contigs using SPAdes version 3.14 ^30^ with parameters “-k 55,77,99,113,127 --careful”. Afterwards, the assembled contigs were reverse complemented, if necessary and rotated to the same start sequence as strain Ames Ancestor. Finally, the contigs were polished once more using Pilon version 1.23 ^31^.

### Bioinformatics analyses

For the long-read assemblies, ribosomal operons were annotated using barrnap version 0.9 (https://github.com/tseemann/barrnap). SNP alleles were searched using USEARCH version 11 ^32^ and the 16S SNP BA/BC probe sequences (see Supplementary Table S2) as an oligo sequence database. To investigate the frequency and distribution of the alleles of 16S rRNA genes in the *B. anthracis* species comprehensively, we downloaded all available short-read Illumina data sets (at the end of 2020) from the NCBI Sequence Read Archive ^33^. These data sets were then assembled using SPAdes v1.14 ^30^ with parameters “-k 55,77,99,113,127 - -careful”. The contigs of the resulting assemblies were extended using tadpole from the BBTools package ^34^ and with parameters “el=1000 er=1000 mode=extend”. Afterwards, blastn ^35^ with parameters “-evalue 1e-10 -word_size 9” was used to align the 23S rRNA sequence against each extended contig end. For each assembly, the number of contigs ending with a 23S rRNA fragment were counted and CanSNPer ^36^ was used to determine the canonical SNPs and likely position in the CanSNP tree. In a next step, kmercountexact from the BBTools package was used with the parameters “fastadump=f mincount=2 k=16” to count all *k*-mers of size 16 from error-corrected reads. From these *k*-mers, the frequencies of the two allelic *k*-mers (sequences of 16 nt used for the dPCR probes) was extracted. kmercountexact also reports a *k*-mer-based coverage estimation of the sequenced reads which is used to filter the assemblies by coverage (min. 20X), number of contigs (max. 200), number of potential rRNAs (>8) and success of CanSNPer prediction. For each remaining assembly, the number of rRNAs carrying the SNP of 16S BA allele was estimated by determine the ratio of the allelic *k*-mers multiplied with the total number of rRNAs, rounded to a whole number. To validate this estimation, we applied the same algorithm to every assembly where both short-and long-reads and/or dPCR results were available and compared the estimated number of BA alleles to the counted number in the long-read assembly or to the measured number from the dPCR experiments. They were consistent across different sequencing coverage, total number of rRNA operons and known BA allele frequencies.

## Data availability

All genomic data generated or analyzed prior or during this study can be accessed via the NCBI BioProject PRJNA695105. Individual accession numbers are listed in Supplementary Tables S3 and S4.

## Acknowledgements

We thank Linda Dobrzykowski and Josua Zinner for technical assistance. For providing bacterial strains we are grateful to Fabian Leendertz, Silke Klee and Roland Grunow (RKI), Wolfgang Beyer (University of Hohenheim) and Paul Keim (Northern Arizona University). We also thank Olfert Landt and his team at TIB Molbiol (Berlin) for technical support in designing LNA-based probes.

This study was supported by funds from the German Federal Ministry of Defense (Sonderforschungsprojekt 36Z1-S-431618 and STAN 48-2009-23).

The research described herein is part of the Medical Biological Defense Research Program of the Bundeswehr Joint Medical Service. Opinions, interpretations, conclusions, and recommendations are those of the authors and are not necessarily endorsed by any governmental agency, department or other institutions.

## Contributions

P.B., F.Z., G.G. and K.S. designed the study and interpreted the results. M.W. contributed the bioinformatics analysis. I.S., F.Z., K.A. and S.M. performed FISH experiments. P.B. and I.S. performed digital PCR experiments. G.G., P.B., F.Z. and K.S. wrote the first draft manuscript and all authors edited the manuscript. The authors dedicate this work to our dear late colleague Karin Aistleitner who left us too soon and unexpectedly.

